# Hypotonic stress induces fast, reversible degradation of the vimentin cytoskeleton via intracellular calcium release

**DOI:** 10.1101/607416

**Authors:** Leiting Pan, Ping Zhang, Fen Hu, Rui Yan, Manni He, Wan Li, Jingjun Xu, Ke Xu

## Abstract

The dynamic response of the cell to osmotic changes is critical to its physiology and has been widely exploited for cell manipulation. Using 3D-STORM super-resolution microscopy, here we examine the hypotonic stress-induced ultrastructural changes of the cytoskeleton of a common fibroblast cell type. Unexpectedly, we observe a fast, yet reversible dissolution of the vimentin intermediate filament system that precedes ultrastructural changes of the supposedly more dynamic actin and tubulin cytoskeletal systems, as well as changes in cell morphology. In combination with calcium imaging and biochemical analysis, we next show that the vimentin-specific fast cytoskeletal degradation under hypotonic stress is due to proteolysis by the calcium-dependent protease calpain. We find the process to be activated by the hypotonic stress-induced calcium release from intracellular stores, and so is efficiently suppressed by inhibiting any part of the IP_3_-Ca^2+^-calpain pathway we establish. Together, our findings highlight an unexpected, fast degradation mechanism for the vimentin cytoskeleton in response to the external stimuli, and point to the significant, yet previously overlooked physiological impacts of hypotonic stress-induced intracellular calcium release on cell ultrastructure and function.

## INTRODUCTION

The intricate inner machinery of the cell depends critically on water homeostasis to regulate intracellular concentrations. Differences in intracellular and external concentrations (osmotic pressures) give rise to osmosis-induced cell volume and behavior changes, which constitute essential processes in physiology and pathology (Ho, 2006; Hoffmann et al., 2009; Jakab et al., 2002). As a cell manipulation tool, hypo-osmotic swelling has been widely utilized for intracellular delivery (Borle and Snowdowne, 1982; Okada and Rechsteiner, 1982; Stewart et al., 2018) and the modulation of membrane tension (Colom et al., 2018; Dai et al., 1998; Groulx et al., 2006). Recent work further established osmotic volume change as a key method to control intracellular protein concentration for studies on protein crowding and protein-protein interactions (Boersma et al., 2015; Sukenik et al., 2017). Consequently, it is of both fundamental and practical importance to understand if osmotic effects would lead to significant intracellular structural changes, and if so, how fast do such changes occur, whether they are reversible and/or suppressible, as well as what mechanisms drive such processes. The cytoskeleton plays essential roles in the cell volume regulation under osmotic stress (Pedersen et al., 2001), a natural consequence given the pivotal role of the cytoskeleton in cell structure and function (Fletcher and Mullins, 2010; Lowery et al., 2015; Pollard and Cooper, 2009).

In particular, the intermediate filament system critically defines the mechanical properties of vertebrate cells (Lowery et al., 2015; Robert et al., 2016). Vimentin is a major component of the intermediate filament system for most cells in culture (Lazarides, 1982); besides supporting the cell shape, it also anchors organelles and impedes intracellular movement by doubling the cytoplasmic shear modulus (Guo et al., 2013; Lowery et al., 2015; Robert et al., 2016). Recent work underscores the dynamic interplays between the vimentin system and the actin and tubulin cytoskeletal systems (Costigliola et al., 2017; Gan et al., 2016; Hookway et al., 2015; Jiu et al., 2015). One emerging theme has been that the vimentin system follows the initial structural cues of the more dynamic actin and tubulin systems, but subsequently helps provide structural persistency through its superior stability and rigidity. Unalterable high stability, however, could pose significant challenges when the cell needs to respond quickly to external stimuli, e.g., that due to osmotic stress.

Using 3D-STORM (three-dimensional stochastic optical reconstruction microscopy) (Huang et al., 2008; Rust et al., 2006) super-resolution microscopy (SRM), here we examine the hypotonic stress-induced ultrastructural changes of the cytoskeleton of a common fibroblast cell type at high spatial resolution. Interestingly, among the three major cytoskeletal systems, we find a quick dissolution of the vimentin intermediate filament system, whereas the actin and tubulin systems are mostly unaffected. We next find the structural changes rapidly recover under normal culturing conditions, with a recovering speed notably faster than that of the cell morphology. In combination with calcium imaging and biochemical analysis, we further establish that the rapid, vimentin-specific cytoskeletal degradation is driven by the calcium binding-mediated activation of the protease calpain through hypotonic stress-induced intracellular calcium release, and so is effectively blocked by corresponding inhibitors and calcium chelators. Together, our findings highlight an unexpected fast degradation mechanism for the vimentin cytoskeleton, and point to the significant, yet previously overlooked physiological impacts of hypotonic stress-induced intracellular calcium release on cell ultrastructure.

## RESULTS

COS-7 cells were cultured in Dulbecco’s Modified Eagle Medium (DMEM) supplemented with 10% FBS following standard tissue-culture protocols. Differential interference contrast (DIC) microscopy (Fig. S1) indicated that after the cells were challenged with the hypotonic stress due to a 50% HBSS (Hank’s balanced salt solution) buffer, no significant changes to cell general morphology was observed at 5 min, although increased membrane ruffling was often noticed at cell edges (Fig. S1A). Meanwhile, cells treated with pure water quickly rounded up within the same time window (Fig. S1B).

To investigate cytoskeletal changes at the ultrastructural level, we fixed the cells for immunofluorescence labeling, and performed 3D-STORM (Huang et al., 2008; Pan et al., 2018; Rust et al., 2006) super-resolution microscopy to achieve ~20 nm spatial resolution optically. Notably, 3D-STORM revealed substantial ultrastructural changes in the vimentin intermediate filaments (Fig. 1AB and Fig. S2A), whereas conventional, diffraction-limited microscopy did not well resolve the filaments (Fig. 1B). For the milder treatment of 50% HBSS buffer, where changes in cell morphology were moderate (Fig. S1A), 3D-STORM showed that the initially long and continuous vimentin filaments, which often ran through the entire cell lengths, all broke down into short segments a few micrometers in length (Fig. 1AB). As a result, the structural network formed by the vimentin filaments (Fig. 1A) disintegrated, presumably negating the force-bearing function of this system (Guo et al., 2013; Lowery et al., 2015; Robert et al., 2016). More dramatically, cells treated with pure water for the same amount of time were characterized by completely dissolved vimentin (Fig. 1AB).

**Fig. 1.**
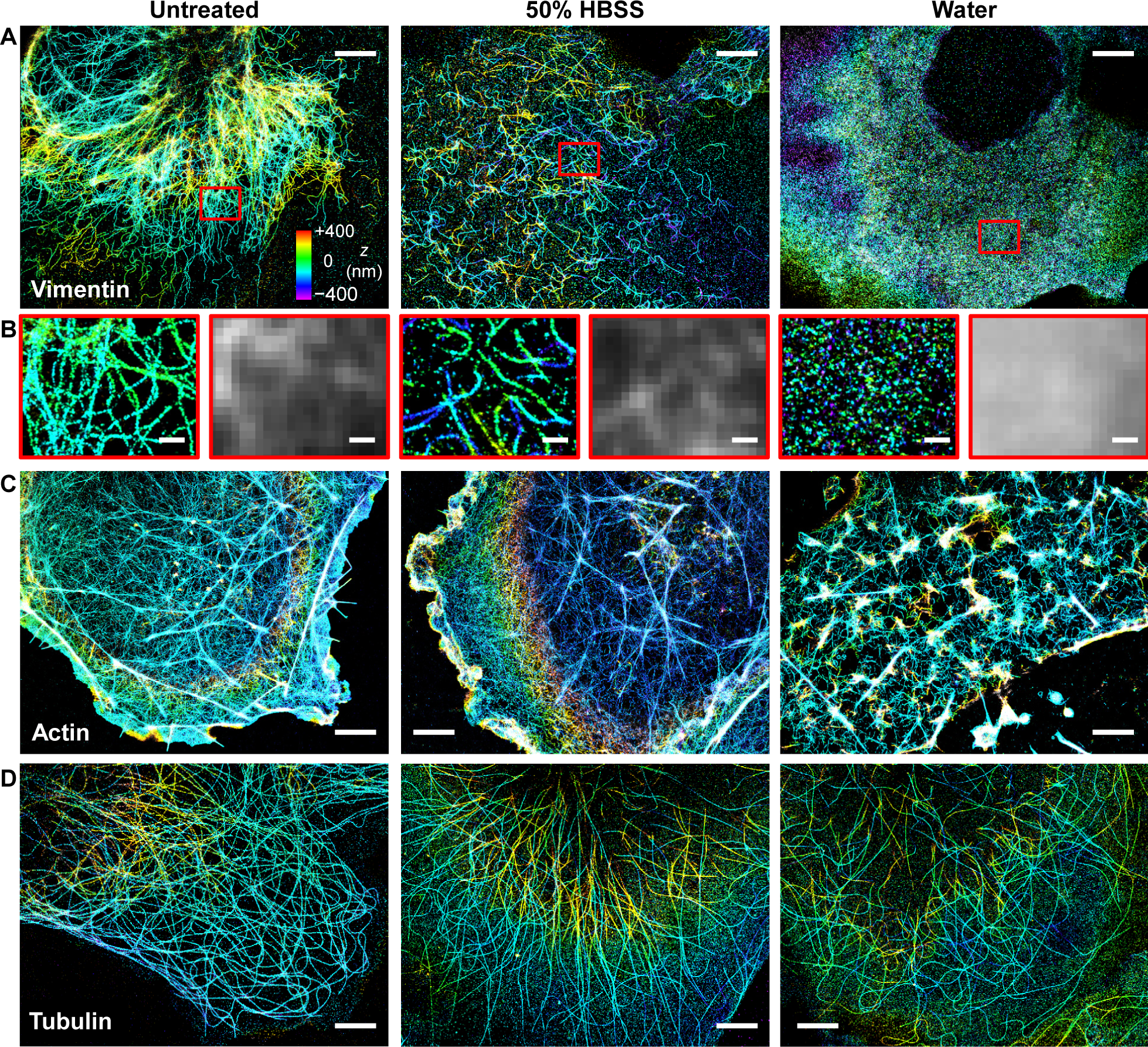
3D-STORM reveals that hypotonic stress leads to fast degradation of the vimentin, but not the actin and tubulin cytoskeletal systems. (A) 3D-STORM images of immunolabeled vimentin for fixed COS-7 cells that were untreated (left), treated by 50% HBSS for 5 min (center), and treated by pure water for 5 min (right). Color represents height *z* (color bar; violet denotes closest to the substrate, and red denotes farthest away). (B) Enlargements of the red boxes in (A), compared to the diffraction-limited epifluorescence images of the same areas. (C) 3D-STORM images of phalloidin-labeled F-actin in fixed COS-7 cells that were untreated (left), treated by 50% HBSS for 5 min (center), and treated by pure water for 5 min (right). (D) 3D-STORM images of immunolabeled alpha-tubulin in fixed COS-7 cells that were untreated (left), treated by 50% HBSS for 5 min (center), and treated by pure water for 5 min (right). Scale bars: 4 µm (A,C,D); 500 nm (B).

In comparison, the actin and tubulin cytoskeletal systems were both found to be much less affected by hypotonic stress (Fig. 1CD and Fig. S2BC). The 50% HBSS treatment did not significantly alter the phalloidin-labeled cortical actin network [Fig. 1C and Fig. S2B; see also more STORM examples of the actin cytoskeleton in (Xu et al., 2012)], except that ruffle-like actin structures were more often observed, in agreement with our DIC results (Fig. S1A). More disrupted cortical actin networks were found for cells treated with pure water (Fig. 1C); however, this disruption appeared not to be due to disassembly of the actin filaments, but instead, was consistent with the reorganization of filaments in response to the high membrane tension (Dai et al., 1998) due to hypotonic swelling. Meanwhile, the ultrastructure of microtubules was largely unaffected by hypotonic stress (Fig. 1D): even for the pure water treatment, microtubules appeared intact, although a somewhat increased staining background was noted, suggesting higher levels of tubulin monomers that did not integrate into the microtubules. Figure S2 provides additional examples of the contrasting ultrastructural changes of the three cytoskeletal systems.

We next investigated if the above ultrastructural changes induced by hypotonic stress were reversible. For cells treated with 50% HBSS, 1 h regrowth in the regular culture medium led to a good recovery of the vimentin cytoskeleton (Fig. 2A), so that the reassembled filaments again ran through entire cell lengths, similar to untreated cells. Cell overall morphology also recovered (Fig. 2A). Interestingly, for cells that had been treated with pure water, 1 h regrowth in the culture medium also led to the full reassembly of long vimentin filaments, even though the general morphology of the cells was still highly abnormal (Fig. 2B). A 24 h regrowth in the culture medium gave satisfactory recovery in terms of both cell morphology and vimentin ultrastructure (Fig. 2C). The actin and tubulin cytoskeletal systems, which were less disrupted by the hypotonic stress in the first place, also recovered after regrowth in the regular culture medium (Fig. S3). Together, our results indicate quick break-down and recovery of the vimentin cytoskeleton under hypotonic stress and regrowth conditions, respectively, and the dynamics of both processes appeared to be substantially faster than that of the cell shape.

**Fig. 2.**
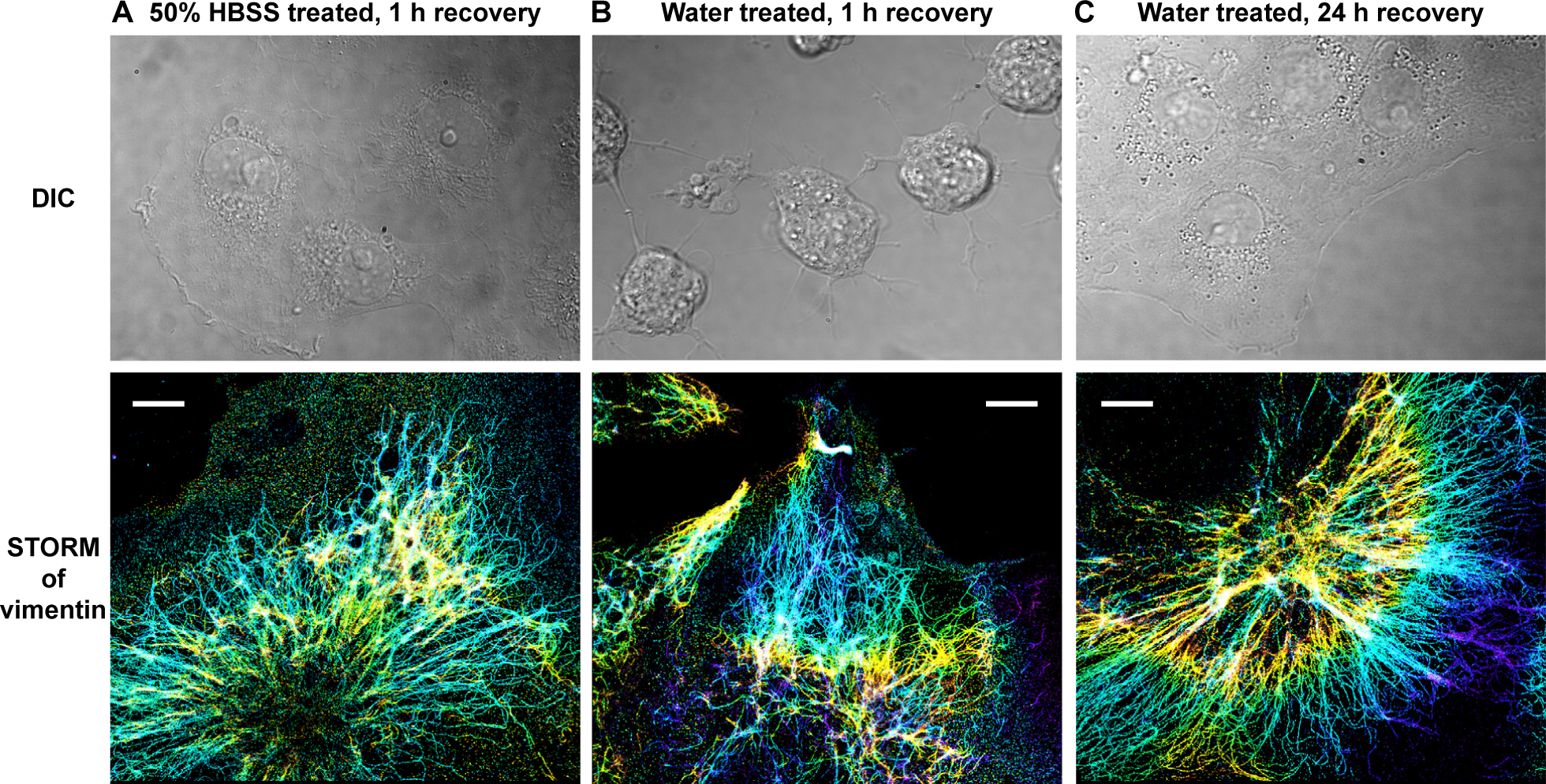
The degraded vimentin network quickly recovers. (A) DIC microscopy (top) and 3D-STORM of vimentin (bottom) for COS-7 cells that had been treated with 50% HBSS for 5 min, and then allowed to recover in the regular culture medium for 1 h. The same color sale as Fig. 1 is used to represent the height *z*. (B) DIC (top) and 3D-STORM of vimentin (bottom) for COS-7 cells that had been treated with pure water for 5 min, and then allowed to recover in the regular culture medium for 1 h. (C) DIC (top) and 3D-STORM of vimentin (bottom) for COS-7 cells that had been treated with pure water for 5 min, and then allowed to recover in the regular culture medium for 24 h. Scale bars: 4 µm.

The striking contrast we observed for how the different cytoskeletal systems responded to the hypotonic stress prompted us to investigate the underlying mechanisms. Paradoxically, whereas actin filaments and microtubules and both known to be structurally highly dynamic and thus prone to dissociation into monomers (Desai and Mitchison, 1997; Pollard and Borisy, 2003), vimentin filaments are often considered highly stable (Lazarides, 1982). Indeed, for structural preservation through chemical fixation, it is well recognized that the proper fixation of actin and microtubules both require strong fixatives like glutaraldehyde and/or specialized cytoskeleton-stabilizing buffers (Small et al., 1999; Svitkina, 2007; Xu et al., 2012), whereas vimentin filaments are readily preserved with common, less potent fixatives as paraformaldehyde (data not shown). Consequently, the observed fast degradation of vimentin filaments, but not actin filaments or microtubules, under hypotonic stress cannot be explained through the intrinsic stabilities of the different filaments.

An important degradation pathway of vimentin is the calcium (Ca^2+^)-dependent proteolysis by calpain family proteins (Goll et al., 2003; Nelson and Traub, 1983). It has been reported that hypotonic stress leads to increased intracellular calcium concentration (Jakab et al., 2002; Missiaen et al., 1996); since the calcium-activation of calpain leads to rapid degradation of vimentin but very slow or no degradation of actin and tubulin (Goll et al., 2003; Nelson and Traub, 1983), a potential differentiation mechanism may thus be established (Fig. 3A).

**Fig. 3.**
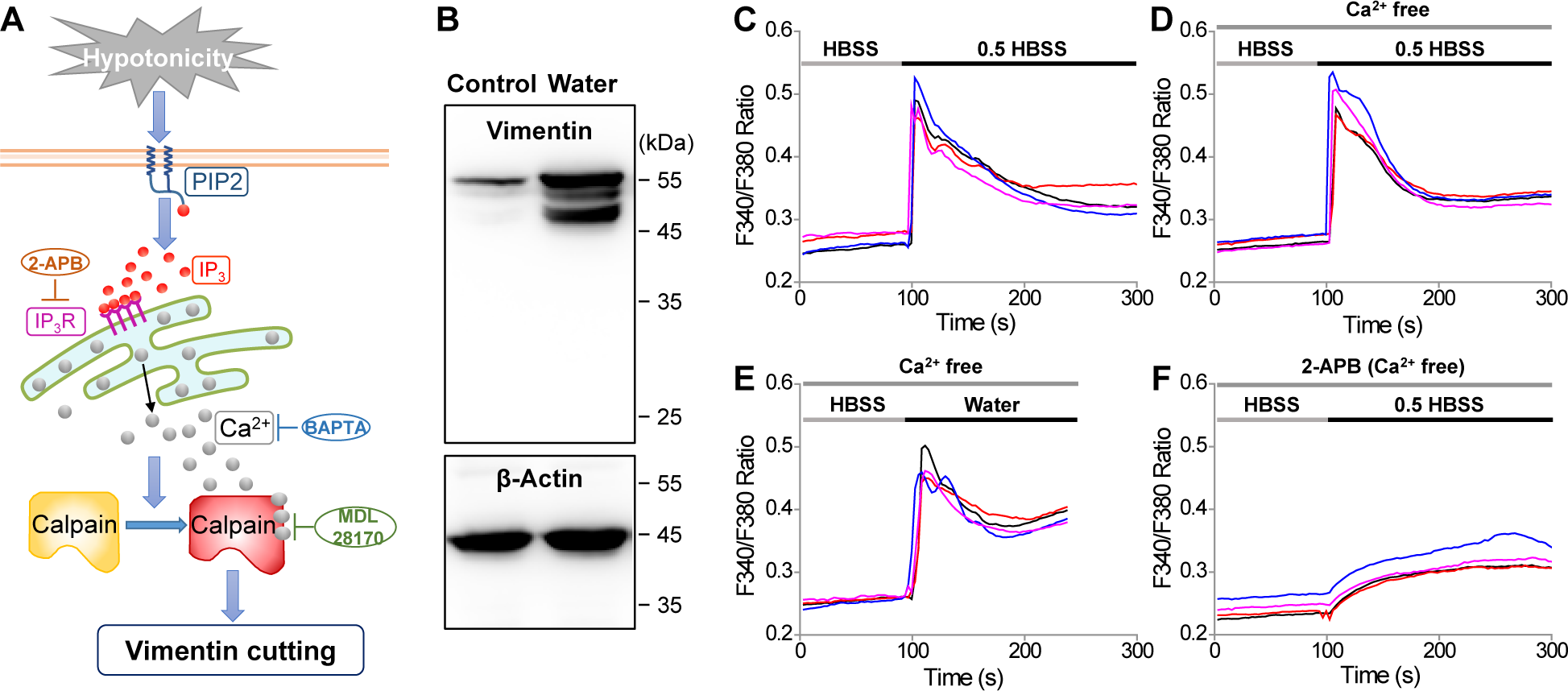
Hypotonic stress induces vimentin cleavage through calcium release from intracellular stores. (A) Schematic for the proposed mechanism of fast vimentin degradation under hypotonic stress. Hypotonic stress leads to an increased level of IP_3_ by hydrolysis of phosphatidylinositol 4,5-bisphosphate (PIP2), which results in calcium release from the ER through the IP_3_ receptor (IP_3_R). Increased cytosolic calcium activates calpain for vimentin cutting. 2-APB is an antagonist for the IP_3_R, BAPTA is a calcium chelator, and MDL 28170 is a calpain inhibitor. (B) Immunoblot analysis of lysates of untreated (control) cells vs. cells that had been treated with water for 5 min. (C-F) Fluorescence readout of cytosolic Ca^2+^, shown as the fura-2 fluorescence intensity ratio at 340 nm vs. 380 nm excitation, for (C) Changing the medium from a regular HBSS buffer to 50% HBSS; (D) Changing the medium from a calcium-free HBSS buffer to a calcium-free 50% HBSS buffer; (E) Changing the medium from a calcium-free HBSS buffer to pure water; (F) Same as (D), but in the presence of 75 µM of 2-APB. For each condition, four typical curves are shown from four different cells.

To test this hypothesis, we performed immunoblot analysis for the lysates of untreated cells and cells that had been subjected to hypotonic stress. A single major band of ~57 kDa was observed for vimentin for the control cells. In contrast, cells that had been treated by water showed two pronounced new bands of lower weights, and intriguingly, all three bands were substantially stronger than the single band observed for the untreated cells (Fig. 3B). The new low-weight bands indicate cleavage products, and the weights we observed (~48 and ~53 kDa) were consistent with calpain-mediated vimentin cleavage (Dourdin et al., 1999; Fischer et al., 1986; Kim et al., 2017). The significantly enhanced band strengths for the water-treated cells further suggest that the vimentin filaments are otherwise highly stable and do not readily dissociate into monomers during the sample preparation, as opposed to the strong cleavage and dissociation after hypotonic treatment, as visualized by SRM above (Fig. 1AB). Moreover, the strength of the cleaved ~48 kDa band was comparable to the original ~57 kDa band, further indicating that a significant fraction of vimentin was cleaved. In comparison, for both samples, β-actin showed a single band of comparable strengths at ~45 kDa (Fig. 3B). Together, our immunoblot analysis suggests that upon hypotonic stress, calpain cleaves vimentin but not actin.

To further examine the possibility of calcium-induced activation of calpain, we next measured cytosolic Ca^2+^ concentrations under hypotonic stress using the ratiometric Ca^2+^ indicator fura-2-acetoxymethyl ester (fura-2/AM). Indeed, challenging the cells with 50% HBSS led to a sudden rise in Ca^2+^ signal (Fig. 3C). To identify whether this rise was due to an influx from the extracellular medium, we repeated the experiment with calcium-free HBSS (Fig. 3D). Comparable rises in Ca^2+^ signal were noted, thus suggesting that the increase in cytosolic Ca^2+^ level was mostly due to calcium release from intracellular stores. In a similar vein, treating the cells with pure water also led to increased cytosolic Ca^2+^ (Fig. 3E).

To further probe the mechanism of this calcium release, we found (Fig. 3F) that the Ca^2+^ increase was suppressed by the application of 2-aminoethoxydiphenyl borate (2-APB), an antagonist for the inositol 1,4,5-trisphosphate (IP_3_) receptor, thus suggesting that the calcium release was due to the IP_3_ signaling pathway (Berridge, 2016) as opposed to passive leakage from the endoplasmic reticulum (ER). Indeed, it has been suggested that hypo-osmotic swelling may result in the hydrolysis of phosphoinositides and an increased intracellular level of IP_3_ (Baquet et al., 1991; Bender et al., 1993; Jakab et al., 2002).

To elucidate whether the above-identified rise in intracellular Ca^2+^ level was indeed the driving force of vimentin degradation, we examined if vimentin degradation could be suppressed by different chemicals that separately block different parts of our proposed IP_3_-Ca^2+^-calpain pathway (Fig. 3A). Indeed, the inclusion of 2-APB, which inhibited intracellular calcium release (Fig. 3F above), substantially suppressed the degradation of vimentin with pure water: 3D-STORM showed well-preserved long vimentin filaments through the entire cell lengths (Fig. 4B and Fig. S4), in stark contrast to the fully dissolved vimentin in control samples treated with water alone (Fig. 4A and Fig. S4). Similar suppression of vimentin degradation was found with the application of BAPTA/AM, a cell-permeable calcium chelator (Fig. 4C and Fig. S4). Finally, MDL 28170, a common calpain inhibitor (Wang and Yuen, 1994), also effectively suppressed vimentin degradation (Fig. 4D and Fig. S4), thus pinpointing the calpain pathway as illustrated in Fig. 3A. Immunoblot analysis of cell lysates also showed that the above inhibitors and calcium chelator to be highly effective in preventing vimentin cleavage and dissociation (Fig. 4E): blocking the upstream calcium signal with 2-APB and BAPTA/AM effectively prevented the generation of cleaved bands, whereas for the calpain inhibitor MDL 28170, a weak ~48 kDa band was observed, indicating substantial, yet incomplete suppression of vimentin degradation.

**Fig. 4.**
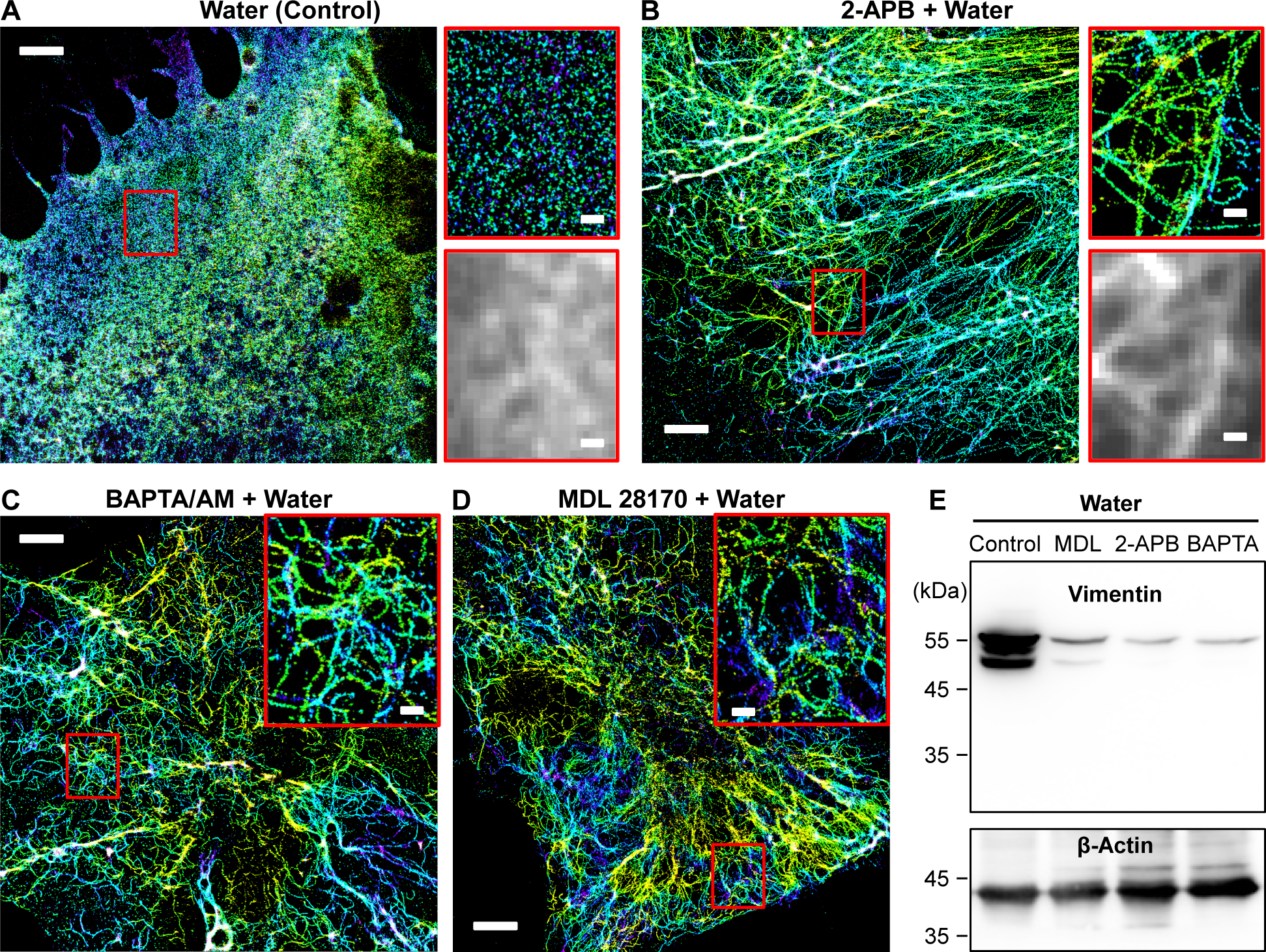
Inhibition of calpain activity or intracellular calcium release preserves the integrity of the vimentin cytoskeleton against hypotonic stress. (A) 3D-STORM image of the immunolabeled vimentin in a control COS-7 cell sample after treatment with pure water for 5 min, together with zoom-in of the red box and the corresponding, diffraction-limited epifluorescence image. The same color sale as Fig. 1 is used to represent the height *z*. (B) 3D-STORM image of a sample treated with water for 5 min in the presence of 75 μM 2-APB, together with zoom-in of the red box and corresponding epifluorescence image. (C) A sample treated with water for 5 min in the presence of 30 μM BAPTA/AM. Inset: zoom-in of the red box. (D) A sample treated with water for 5 min in the presence of 30 μM MDL 28170. Inset: zoom-in of the red box. Scale bars: 4 µm (main figures) and 500 nm (zoom-ins). (E) Immunoblot analysis of cell lysates corresponding to the conditions in (A)-(D).

## DISCUSSION

Hypotonic processes are essential both for their key roles in cell physiology and pathology and for their wide use as cell manipulation tools. Through 3D-STORM SRM, we showed that hypotonic stress induced rapid degradation of the vimentin cytoskeleton, whereas the actin and microtubule cytoskeletal systems were much less affected (Fig. 1). Our finding that disruptions to the cell ultrastructure were fully recoverable under normal cell culture conditions (Fig. 2) suggests that the use of hypo-osmotic swelling for intracellular delivery (Borle and Snowdowne, 1982; Okada and Rechsteiner, 1982; Stewart et al., 2018) should be acceptable as long as enough recovery time is provided. Conversely, for experiments on the use of hypo-osmotic swelling to modulate membrane tension (Colom et al., 2018; Dai et al., 1998; Groulx et al., 2006) and protein concentration (Boersma et al., 2015; Sukenik et al., 2017), it may be necessary to account for the additional intracellular structural changes when interpreting data, as opposed to considering changes in cell geometry alone. Fortunately, in this work we also identified effective, facile means to suppress such structural changes (Fig. 4). It would thus be helpful to reexamine if different results might be obtained when the above structural changes are suppressed.

Our results highlight the contrasting responses of different cytoskeletal systems to external stimuli. Unexpectedly, while the actin and tubulin systems are often considered as being more structurally dynamic, we found that under hypotonic stress, fast degradation was specific to the vimentin intermediate filaments (Fig. 1). This counterintuitive result, which we later found to be due to a unique IP_3_-Ca^2+^-calpain pathway, helps resolve the above-discussed conundrum of how the otherwise highly stable and rigid vimentin filaments could respond quickly to the hypotonic stress, so as to allow the cell to rearrange morphologically and internally.

Our work further demonstrates significant, yet previously overlooked physiological impacts of hypotonic stress-induced intracellular calcium release, as well as its downstream effects through calpain activation (Fig. 3). Notably, we showed that the hypotonic stress-induced calpain activation could be effectively suppressed by blocking any part of the IP_3_-Ca^2+^-calpain pathway (Fig. 4). While these results already suggest hypotonic stress could be utilized as a facile, yet potent tool to induce intracellular calcium release and calpain activation, it would be further interesting to ask what other effects may also be triggered by the same pathway, which should also be similarly suppressible by the same inhibitors. Besides intermediate filaments, in neuronal cells and blood cells, members of the spectrin-based cortical membrane cytoskeleton (Bennett and Gilligan, 1993; Pan et al., 2018; Xu et al., 2013) are also important substrates of calpain (Czogalla and Sikorski, 2005; Goll et al., 2003). Indeed, for the neuron axon initial segment, it has been shown that injury-induced increase in cytosolic calcium activates calpain toward the rapid degradation of βIV spectrin and ankyrin G, hence an important mechanism of neuronal injury (Schafer et al., 2009). Meanwhile, in the ocular lens, Ca^2+^ elevation caused by altered ubiquitin activates calpain toward the degradation of vimentin, fodrin (αII spectrin), and other proteins, hence developmental defects and cataract (Liu et al., 2015). Our finding of hypotonic stress-induced Ca^2+^ elevation and calpain activation may thus carry broad implications for the structure and function of cells under related physiological and pathological conditions.

## EXPERIMENTAL PROCEDURES

### Regents

Dulbecco’s Modified Eagle Medium (DMEM) and Hank’s Balanced Salt Solution (HBSS) were from Gibco. EM-grade paraformaldehyde (15714) and glutaraldehyde (16365) were purchased from Electron Microscopy Sciences (Hatfield, PA, USA). Fura-2/AM (F0888), MDL 28170 (M6690), 2-Aminoethyl diphenylborinate (2-APB) (D9754), cysteamine (30070), glucose oxidase (G2133), catalase (C30), MES (69892), bovine serum albumin (BSA) (A3059), and other general reagents were obtained from Sigma-Aldrich (St Louis, MO, USA). BAPTA/AM (B6769), and Alexa Fluor 647-conjugated phalloidin (A22287) were from Invitrogen (Carlsbad, CA, USA). Primary antibodies used: Vimentin, chicken polyclonal, Millipore AB5733; Tubulin, mouse monoclonal, Abcam ab7291. Alexa Fluor 647-conjugated secondary antibodies (A31571 and A21449, Invitrogen) were used for single-color STORM.

### Cell culture and treatments

COS-7 cells were cultured in DMEM supplemented with 10% FBS in a humidified CO_2_ incubator with 5% CO_2_ at 37°C, following standard tissue-culture protocols. Hypotonic stress was applied by replacing the culture medium with a mixed 1:1 HBSS:water solution (50% HBSS) or pure water at room temperature. For the recovery experiments, the 50% hypo-osmotic solution or pure water was replaced by the original culture medium, and the sample was kept as above in the incubator for an additional indicated time course. To assess the effects of inhibitors, cells were first pre-incubated with 75 μM 2-APB, 30 μM MDL 28170 for 15 min in HBSS or 30 μM BAPTA/AM in HBSS for 30 min, and the sample was treated with pure water containing the same inhibitors at the same concentrations for 5 min.

### Measurement of cytosolic Ca^2+^ concentrations ([Ca^2+^]_c_)

[Ca^2+^]_c_ was measured by a calcium imaging system built on an inverted fluorescent microscope (Olympus IX51, Japan) using the ratiometric Ca^2+^ indicator fura-2/AM. Cells were first loaded with 5 μM fura-2/AM in HBSS for 1 h at room temperature in the dark. After a gentle washing step, cells were bathed in a fresh HBSS (or calcium-free HBSS, as indicated in the text) solution for [Ca^2+^]_c_ measurement. The cells were alternately excited by a Xenon lamp at 340 and 387 nm using a motorized filter wheel (Lambda 10-2, Sutter Instrument, Novato, CA, USA). Fluorescence images (filtered at 515 nm ± 25 nm) were acquired by a CCD camera (CoolSNAP fx-M, Roper Scientific, Tucson, AZ, USA) and analyzed with MetaFluor (Universal Imaging, West Chester, PA, USA). [Ca^2+^]_c_ was represented by the ratio of fluorescence intensity excited at 340 nm / excited at 380 nm (F340/F380).

### Immunoblot analysis of cell lysates

Cells were seeded into 6-well plates at 1×10^6^/well and incubated overnight in DMEM with 10% FBS at 37°C. Total protein lysates were isolated by RIPA (150 μL/well, R0020, Solarbio, China). The concentration of protein was determined using a BCA assay kit (P0012S, Beyotime, China). Afterward, aliquots (20 μg/lane) were separated by 10% SDS-PAGE and transferred to a nitrocellulose membrane, blocked with 5% BSA at room temperature for 1 h, immunoblotted with rabbit monoclonal antibody against vimentin (1:1000, #5741; Cell Signaling Technology, Danvers, MA, USA) as well as mouse monoclonal antibody against β-actin (1:20000, 60008-1-Ig; Proteintech, Rosemont, IL, USA) overnight at 4°C, followed by incubation with anti-rabbit/mouse horseradish peroxidase-conjugated secondary antibody (1:1000; A0208/A0216, Beyotime, China) respectively. Finally, an ECL detection reagent (B500012, Proteintech, USA) was used for visualization in a Tanon 5200 MultiImage System.

### Cell fixation and immunofluorescence

Cells were seeded on 12-mm glass coverslips in a 24-well plate at ~2×10^4^ cells per well, and cultured for 12 h. For STORM of actin filaments, a previously established fixation protocol was employed (Small et al., 1999; Svitkina, 2007; Xu et al., 2012): The samples were initially fixed and extracted for 1 min using a solution of 0.3% (v/v) glutaraldehyde and 0.25% (v/v) Triton X-100 in cytoskeleton buffer (CB, 10 mM MES, pH 6.1, 150 mM NaCl, 5 mM EGTA, 5 mM glucose and 5 mM MgCl_2_), and then post-fixed for 15 min in 2% (v/v) glutaraldehyde in CB, and reduced with a freshly prepared 0.1% sodium borohydride solution in PBS. Alexa Fluor 647-conjugated phalloidin was applied at a concentration of ~0.4 µM for 1 h. The sample was briefly washed 2-3 times with PBS, and then immediately mounted for imaging. For imaging of other targets, samples were fixed with 3% (w/v) paraformaldehyde and 0.1% (w/v) glutaraldehyde in phosphate buffered saline (PBS) for 20 min. After reduction with a freshly prepared 0.1% sodium borohydride solution in PBS for 5 min, the samples were permeabilized and blocked in a blocking buffer (3% w/v BSA, 0.5% v/v Triton X-100 in PBS) for 20 min. Afterward, the cells were incubated with the primary antibody (described above) in blocking buffer for 1 h. After washing in washing buffer (0.2% w/v BSA and 0.1% v/v Triton X-100 in PBS) for three times, the cells were incubated with the secondary antibody for 1 h at room temperature. Then, the samples were washed 3 times with washing buffer before mounted for imaging.

### Super-resolution microscopy

After washing with PBS, the samples were mounted on glass slides with a standard STORM imaging buffer consisting of 5% (w/v) glucose, 100 mM cysteamine, 0.8 mg/mL glucose oxidase, and 40 µg/mL catalase in Tris-HCl (pH 7.5) (Huang et al., 2008; Rust et al., 2006). Then, data were collected by 3D-STORM (Huang et al., 2008; Rust et al., 2006) carried out on a homebuilt setup based on a modified Nikon Eclipse Ti-E inverted optical microscope using an oil-immersion objective (Nikon CFI Plan Apochromat λ, 100×, numerical aperture = 1.45), as described in (Wojcik et al., 2015). Lasers at 405 and 647 nm were introduced into the cell sample through the back focal plane of the objective, and shifted towards the edge of the objective to illuminate ~1 µm within the glass-water interface. A strong (~2 kW cm^-2^) excitation laser of 647 nm photoswitched most of the labeled dye molecules into a dark state, while also exciting fluorescence from the remaining, sparsely distributed emitting dye molecules for single-molecule localization. A weak (typical range 0-1 W cm^-2^) 405-nm laser was used concurrently with the 647-nm laser to reactivate fluorophores into the emitting state, so that at any given instant, only a small, optically resolvable fraction of fluorophores was in the emitting state. A cylindrical lens was put into the imaging path to introduce astigmatism to encode the depth (z) position into the ellipticity of the single-molecule images (Huang et al., 2008). The Andor iXon Ultra 897 EMCCD concurrently recorded images at 110 frames per second for a frame size of 256×256 pixels, and typically recorded ~50,000 frames for each experiment.

## SUPPLEMENTAL INFORMATION

Supplemental Information includes four figures.

## ACKNOWLEDGMENTS

This work was supported by the National Natural Science Foundation of China (no. 11874231, 11574165 and 31801134), Tianjin Natural Science Foundation (no. 18JCQNJC02000), the PCSIRT (no. IRT_13R29), the 111 Project (no. B07013), and the Pew Biomedical Scholars Award. K.X. is a Chan Zuckerberg Biohub investigator.

## Supplemental Figures

**Figure S1.**
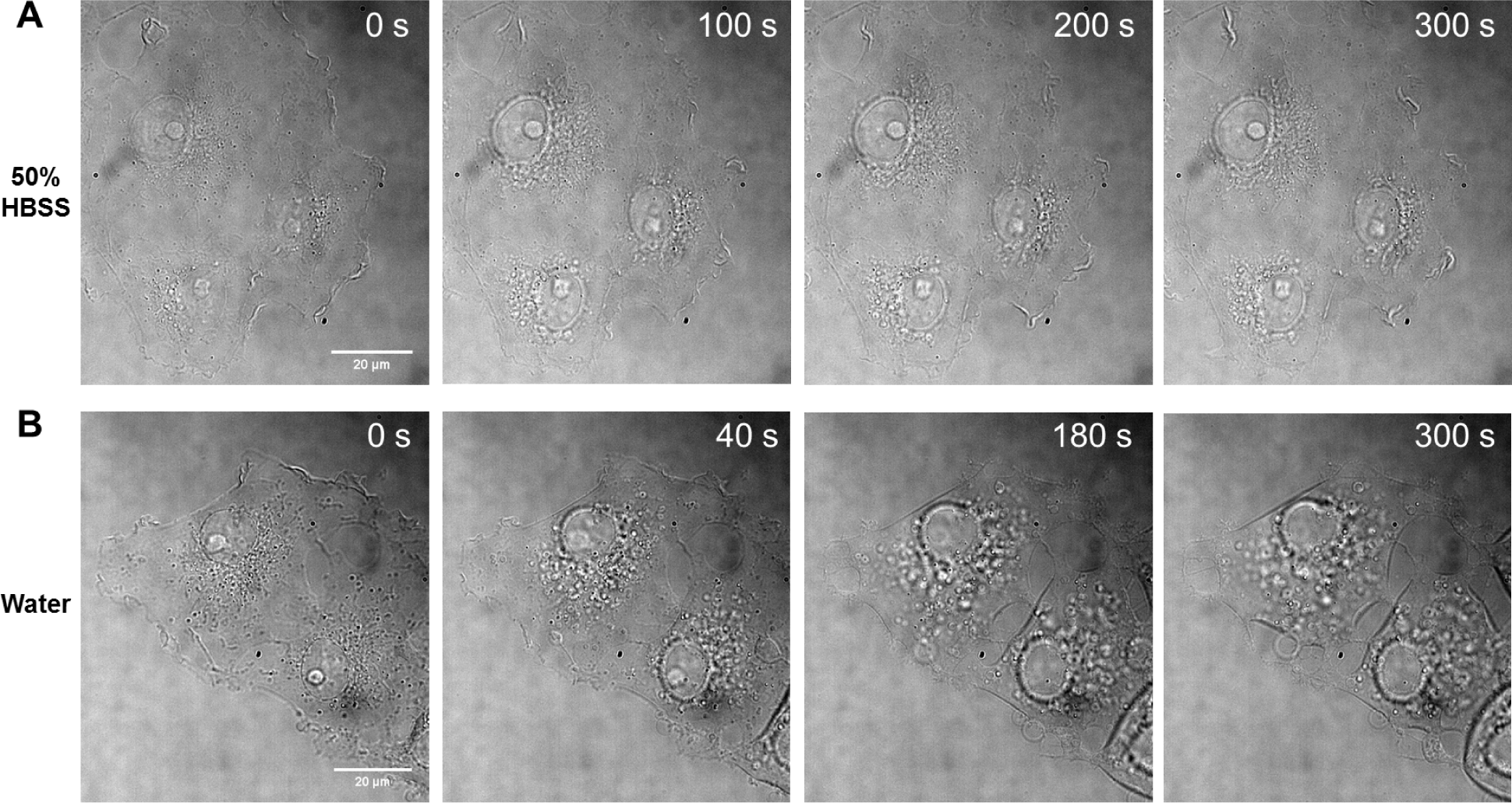
DIC microscopy of cell morphology under hypotonic stress. (A) Image sequences for COS-7 cells treated with 50% HBSS. (B) Image sequences for COS-7 cells treated with pure water.

**Figure S2.**
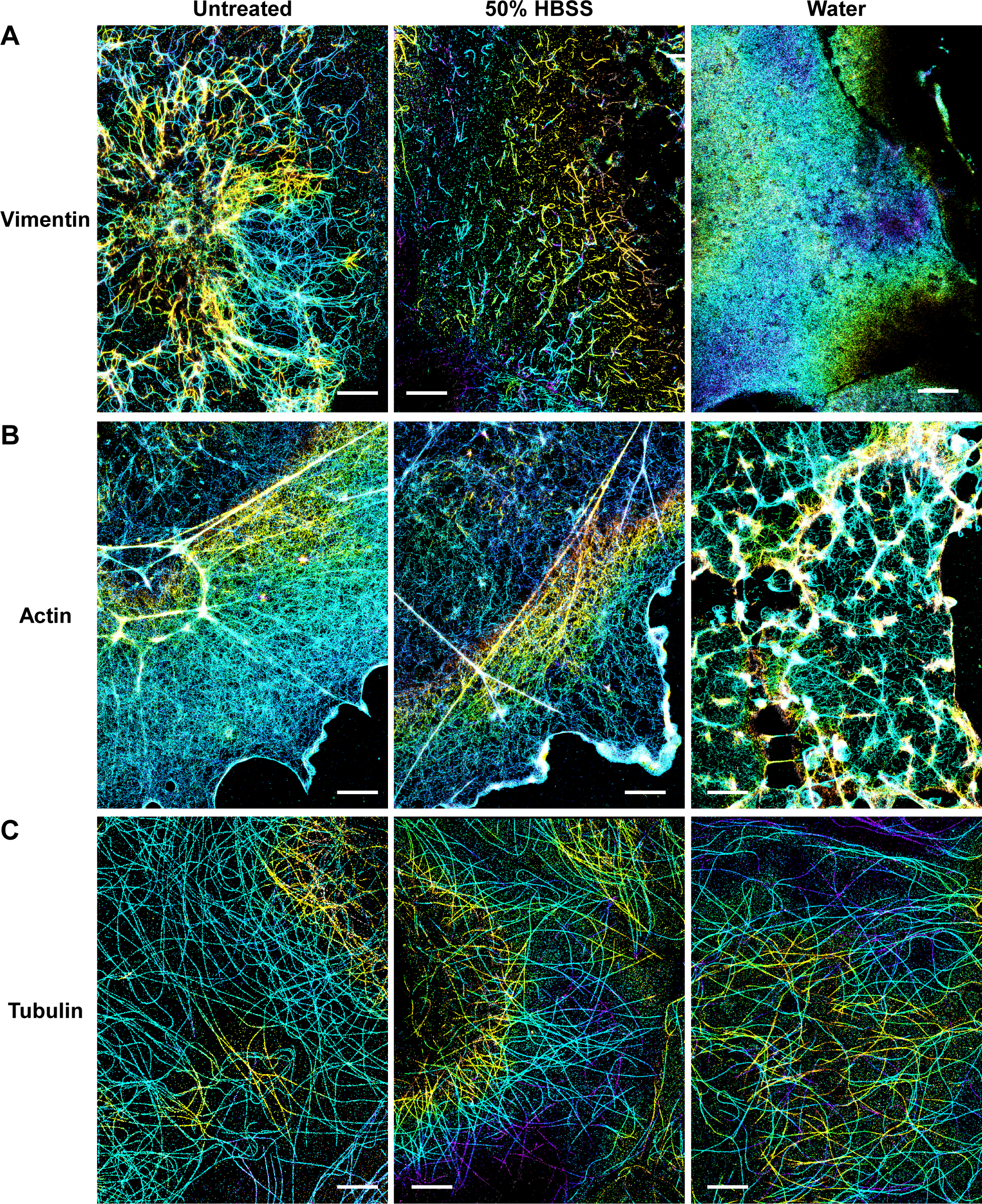
Additional 3D-STORM images of the three cytoskeletal systems under hypotonic stress. (A) Immunolabeled vimentin in COS-7 cells untreated (left), treated by 50% HBSS for 5 min (center), and treated by pure water for 5 min (right). (B) Phalloidin-labeled F-actin. (C) Immunolabeled alpha-tubulin. Scale bars: 4 µm.

**Figure S3.**
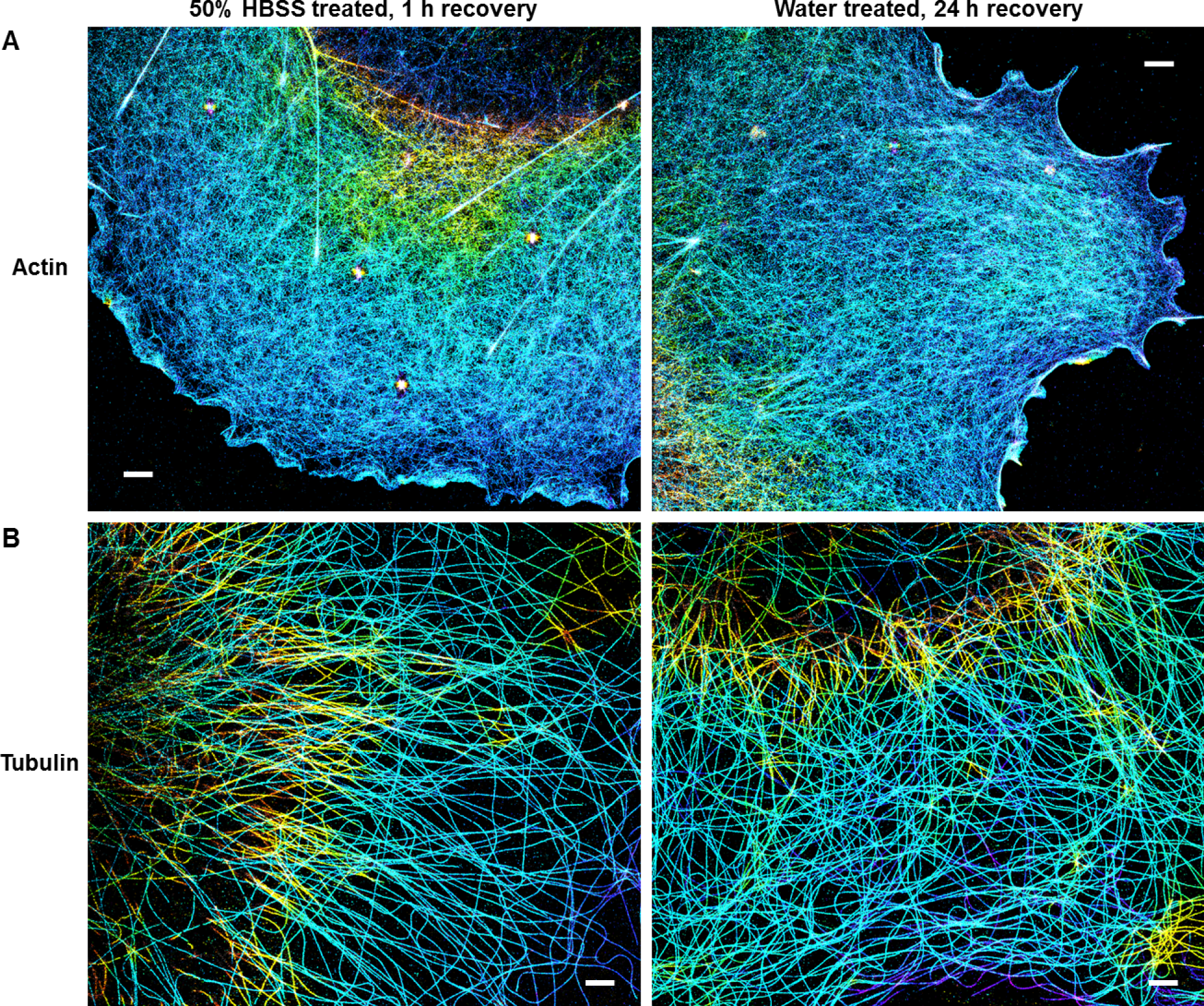
Recovery of the actin and tubulin cytoskeletal systems. (A) 3D-STORM images of phalloidin-labeled F-actin in COS-7 cells that had been treated with 50% HBSS for 5 min and then allowed to recover in the regular culture medium for 1 h (left), and that had been treated with pure water for 5 min, and then allowed to recover in the regular culture medium for 24 h (right). (B) 3D-STORM images of immunolabeled alpha-tubulin for cells under the same conditions. Scale bars: 2 µm.

**Figure S4.**
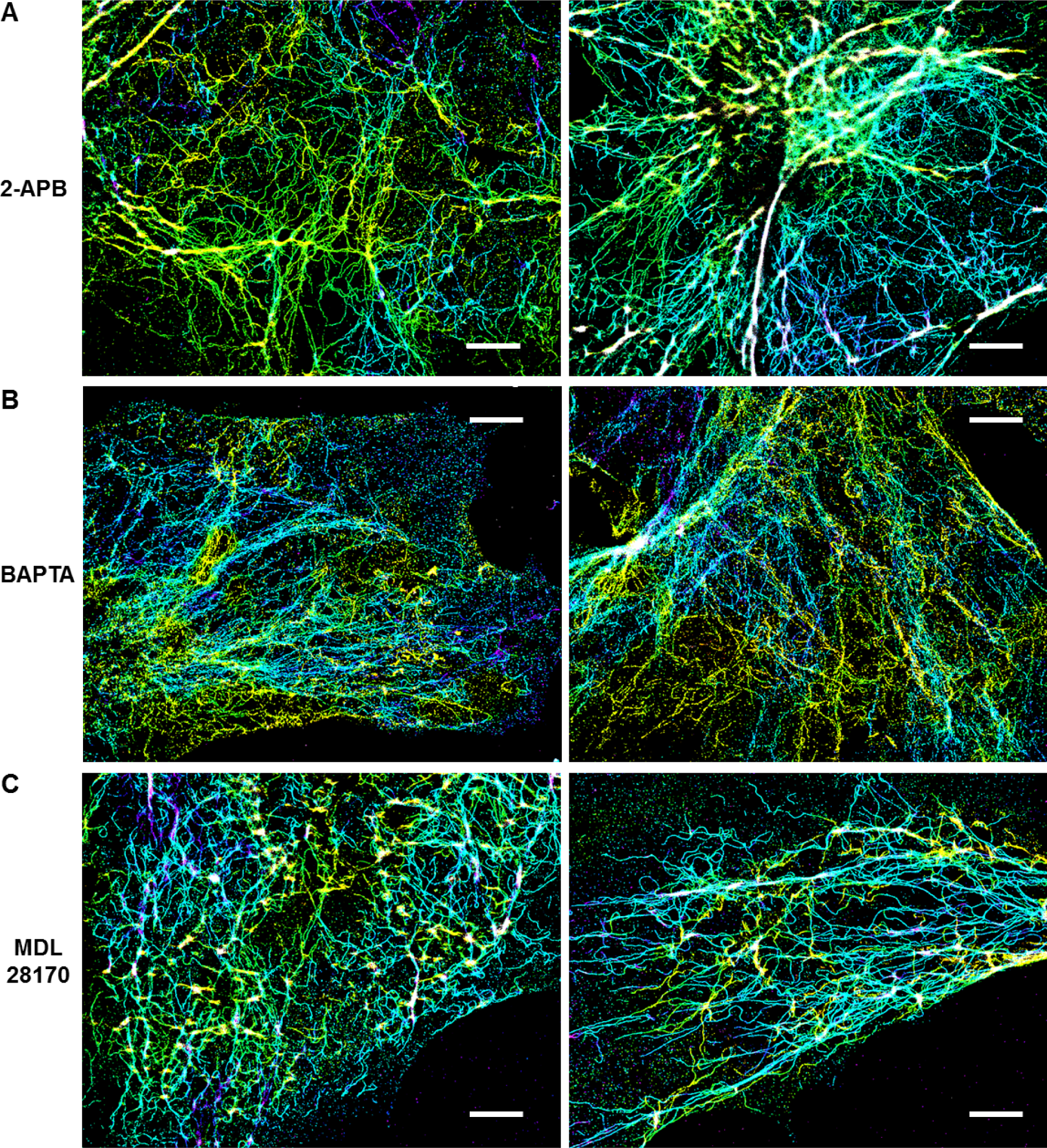
Additional examples of preserving the vimentin cytoskeleton against hypotonic stress through blocking the IP_3_-Ca^2+^-calpain pathway. (A) 3D-STORM images of immunolabeled vimentin in COS-7 cells treated with water for 5 min in the presence of 75 μM 2-APB. (B) Cells treated with water for 5 min in the presence of 30 μM BAPTA/AM. (C) Cells treated with water for 5 min in the presence of 30 μM MDL 28170. Scale bars: 4 µm.

